# The role of glycaemic, lipid, blood pressure and obesity risk factors as mediators of the effect of height on Coronary Artery Disease and Type 2 Diabetes Mellitus: A Mendelian Randomisation Study

**DOI:** 10.1101/468918

**Authors:** Eirini Marouli, Del Greco M. Fabiola, Christina Astley, Jian Yang, Shafqat Ahmad, Sonja Berndt, Mark J. Caulfield, Evangelos Evangelou, Barbara McKnight, Carolina Medina-Gomez, Jana V. van Vliet-Ostaptchouk, Helen R. Warren, Zhihong Zhu, GIANT Consortium

**Affiliations:** William Harvey Research Institute, Barts and The London School of Medicine and Dentistry, Queen Mary University of London, London, UK; Centre for Genomic Health, Life Sciences, Queen Mary University of London, London, UK; Institute for Biomedicine, Eurac Research, Affiliated Institute of the University of Lubeck, Bolzano, Italy; Boston Children’s Hospital, MA, USA; Broad Institute of Harvard and MIT, Cambridge, MA, USA; Institute for Molecular Bioscience, University of Queensland, Brisbane, Queensland, Australia; Queensland Brain Institute, The University of Queensland, Brisbane, Queensland, Australia; Department of Nutrition, Harvard T.H. Chan School of Public Health, Harvard University, Boston,MA, USA; Division of Preventive Medicine, Harvard Medical School, Department of Medicine, Brigham and Women’s Hospital, Boston, MA, USA; Department of Medical Sciences, Molecular Epidemiology, Uppsala University, Uppsala, Uppsala, Sweden; Division of Cancer Epidemiology and Genetics, National Cancer Institute, National Institutes of Health, Department of Health and Human Services, Bethesda, Maryland, USA; National Institute for Health Research, Barts Cardiovascular Biomedical Research Center, Queen Mary University of London, London, UK; Department of Epidemiology and Biostatistics, School of Public Health, Imperial College London, London, UK; Department of Hygiene and Epidemiology, University of Ioannina Medical School, Ioannina, Greece; Department of Biostatistics, University of Washington, Seattle, WA, USA; Department of Internal Medicine, Erasmus Medical Center, Rotterdam, The Netherlands; Department of Epidemiology, Erasmus Medical Center, Rotterdam, The Netherlands; Department of Endocrinology, University of Groningen, University Medical Center Groningen, Groningen, The Netherlands; The Charles Bronfman Institute for Personalized Medicine, Icahn School of Medicine at Mount Sinai, New York, NY, USA; Institute of Social and Preventive Medicine, Lausanne University Hospital, Lausanne, Switzerland; Swiss Institute of Bioinformatics, Lausanne, Switzerland; Princess Al-Jawhara Al-Brahim Centre of Excellence in Research of Hereditary Disorders (PACER-HD), King Abdulaziz University, Jeddah, Saudi Arabia

## Abstract

We investigated by Mendelian Randomisation (MR) analyses the causal effect of adult height on coronary artery disease (CAD) (23,755 cases, 425,339 controls) and type 2 diabetes (T2D) (29,427 cases, 416,908 controls) in UK Biobank. We then examined whether such effects are mediated via known cardiometabolic risk factors including body mass index (BMI), glycaemic traits, lipid levels and blood pressure. We further cross-checked our findings by two-sample MR and Multivariate MR analyses with publicly available summary statistics data. One standard deviation (SD) higher genetically determined height (~6.5 cm) was causally associated with a 14% decrease in CAD risk (OR= 0.86, 95% CI= 0.79-0.94). This causal association remained significant after performing sensitivity analyses relaxing pleiotropy assumptions. The causal effect of height on CAD risk appeared to be independent of risk factors such as lipid levels, blood pressure and BMI; association was reduced by only 1-3% after adjustment for potential mediators. We observed no direct causal effect of height on T2D risk, once its effect on BMI was taken into account (OR= 0.99, 95% CI= 0.96-1.02), suggesting that BMI acts as a mediator.

---

Evidence from observational studies suggests that height is associated with different disease outcomes ^1, 2, 3, 4^ For example, a decrease of 1 SD in genetically determined height (~6.5 cm) has been associated with a 13% higher risk of coronary artery disease (CAD)^4^.

In situations where randomized trials are inappropriate or impossible, Mendelian Randomisation (MR) provides a good alternative to study the causal relationship between a trait and a disease outcome. MR which is an instrumental variable (IV) – based method to infer causality in observational studies ^5^, can, therefore, be applied to investigate any causal effect that adult height may have on cardiometabolic outcomes, and provide some insight about potential mechanisms. MR offers major advantages; for example, germ-line genetic variants are assorted during formation of gametes prior to conception and are not confounded by lifestyle or environmental factors in ethnically homogeneous samples of unrelated individuals. Thus, it becomes possible to investigate how height variants may affect cardiometabolic risk and whether this effect is direct or mediated through other biological pathways.

MR relies on the availability of genetic variants robustly associated with height. So far, large-scale meta-analyses of genome-wide association studies (GWAS) have identified circa 600 loci associated with adult height^6, 7^ harbouring over 960 independent associations. In our latest study, height-increasing alleles at all 606 height-associated variants (Exome Chip data) were enriched for nominally significant protective effect on several cardiometabolic traits (TC; *P*binomial = 4.4 ×10^−8^), triglyceride (TG; *P*binomial =8.9×10^−7^) and coronary artery disease (CAD; *P*binomial =6.0×10^−10^).

Besides CAD, greater adult stature has also been reported to be associated with lower risk of type 2 diabetes (T2D) ^8^. Adult stature is the result of bone elongation and bone serves as a scaffold for other organs and is an endocrine organ involved in the regulation of glucose and energy metabolism, consequently, hormones implicated in bone re-modelling may affect risk of cardiometabolic disease ^9^.

To test whether height is causally related to cardiometabolic disease (CAD and T2D), including traditional risk factors, we undertook MR analyses in UK Biobank (UKBB)^10^ We performed instrumental variable analysis using individual data and 829 of the previously established height-associated SNPs, which explain around 30% of height variation ^6, 7^. We also applied two-sample MR analyses, including: Inverse-variance-weighted – (IVW); MR-Egger (Egger); Weighted Median (WM) and Mode-based estimate (MBE) methods ^11, 12^, using publically available summary level data from GIANT (anthropometric traits), CARDIOGRAMplusC4D (CAD), DIAGRAM (T2D), MAGIC (glycaemic traits), GLGC (lipid traits) and ICBP (BP). In addition, we investigated whether the effect of height on cardiometabolic disease risk (CAD and T2D) was mediated by glycaemic (glucose, insulin, glycated hemoglobin (HBA1c), 2 hours postprandial glucose-2hGlu), blood pressure (Systolic Blood Pressure (SBP), Diastolic Blood Pressure (DBP), Pulse Pressure (PP)), obesity traits (BMI) and lipid (Total Cholesterol (TC), Low Density Lipoprotein (LDL), High Density Lipoprotein (HDL), Triglycerides (TG)) traits, using both, summary data and individual level data from UBBB where available. A flowchart of all analyses undertaken is presented in Figure 1.

**Figure 1.**
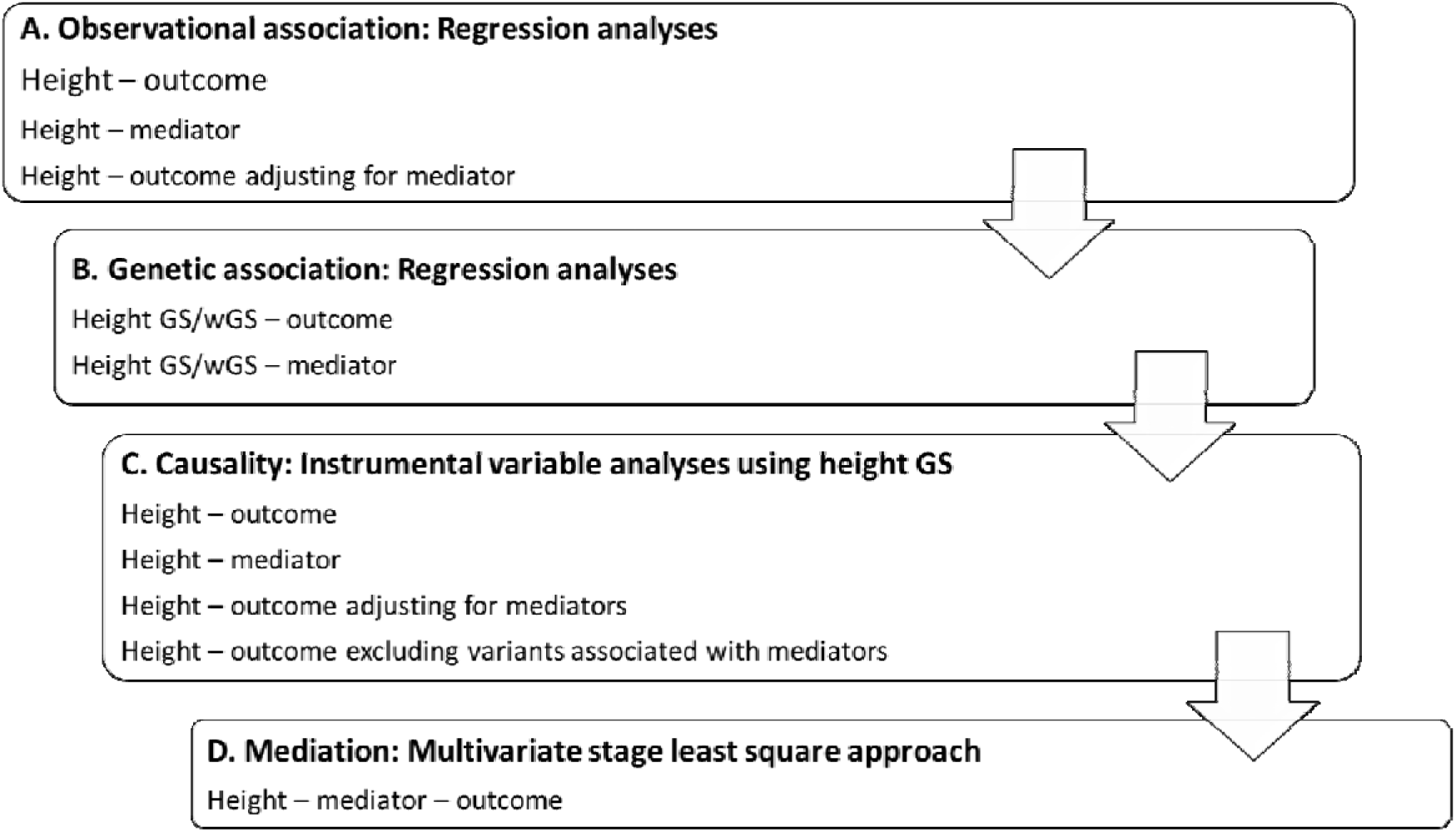
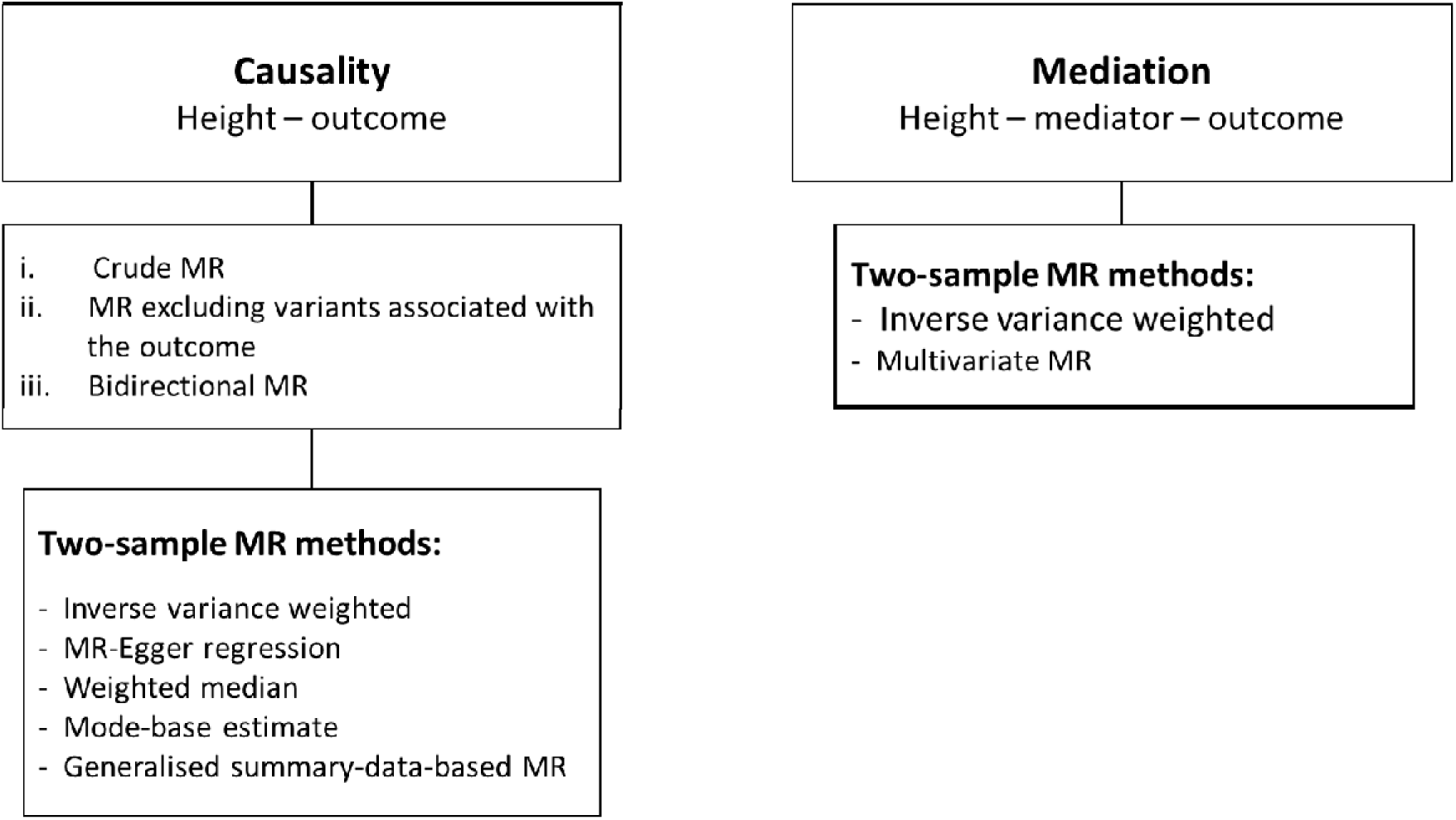
Flowchart of the study design. A: Using individual level data from UK Biobank B: Using summary data

## Results

To test whether genetically determined height is related to cardiometabolic disease phenotypes, independently of traditional risk factors, we undertook MR analyses in UKBB including 449,094 unrelated British participants with both phenotypic and genetic data (STable 1). Mean height was 168.53 cm (range: 125-209 cm); 23,755 individuals had CAD and 29,427 had T2D (see Methods for inclusion criteria). To perform MR analyses, we constructed an unweighted and a weighted genetic score (GS and wGS respectively) using 829 SNPs associated with adult height. The two scores were normally distributed in UKBB and robustly associated with height, as expected, in this cohort (SFigures 1-2).

### Measured and genetically determined height associations

We initially tested for association between measured adult height and cardiometabolic diseases (CAD and T2D) (STable 2a). A 1 SD increase in height was associated with an OR of 0.82 (95% CI= 0.81 to 0.83) and 0.89 (95% CI= 0.87 to 0.90) for risk of CAD and T2D, respectively, consistent with previously reported associations ^3, 4^. We also tested the effect of height on cardiometabolic disease by taking into account risk factors, but the observed effects were not affected (STable 2b). A higher GS was associated with a protective effect on CAD risk (OR = 0.84, 95% CI= 0.81, 0.87) and T2D risk (OR = 0.92, 95% CI= 0.89 to 0.95) (STable 3).

### Mendelian Randomisation analyses

Having established observational and genetic associations between adult height and CAD and T2D risk respectively, we set to perform MR analyses to further investigate whether this relationship is causal or not.

### Instrumental variable analysis in the UKBB

Two-stage analyses for CAD and T2D events in UKBB, using either the unweighted or the weighted GS (STable 4a), showed in all instances an inverse association. For the GS, a 1 SD higher height was associated with an OR of 0.78 (95% CI= 0.74 to 0.82) for CAD and an OR of 0.89 (95% CI= 0.85 to 0.93) for T2D. A similar effect was observed when using the wGS instrument (OR of 0.81 (95% CI= 0.78 to 0.85) for CAD and an OR of 0.92 (95% CI= 0.88 to 0.95) for T2D) (STable 4a, Figure 2).

**Figure 2:**
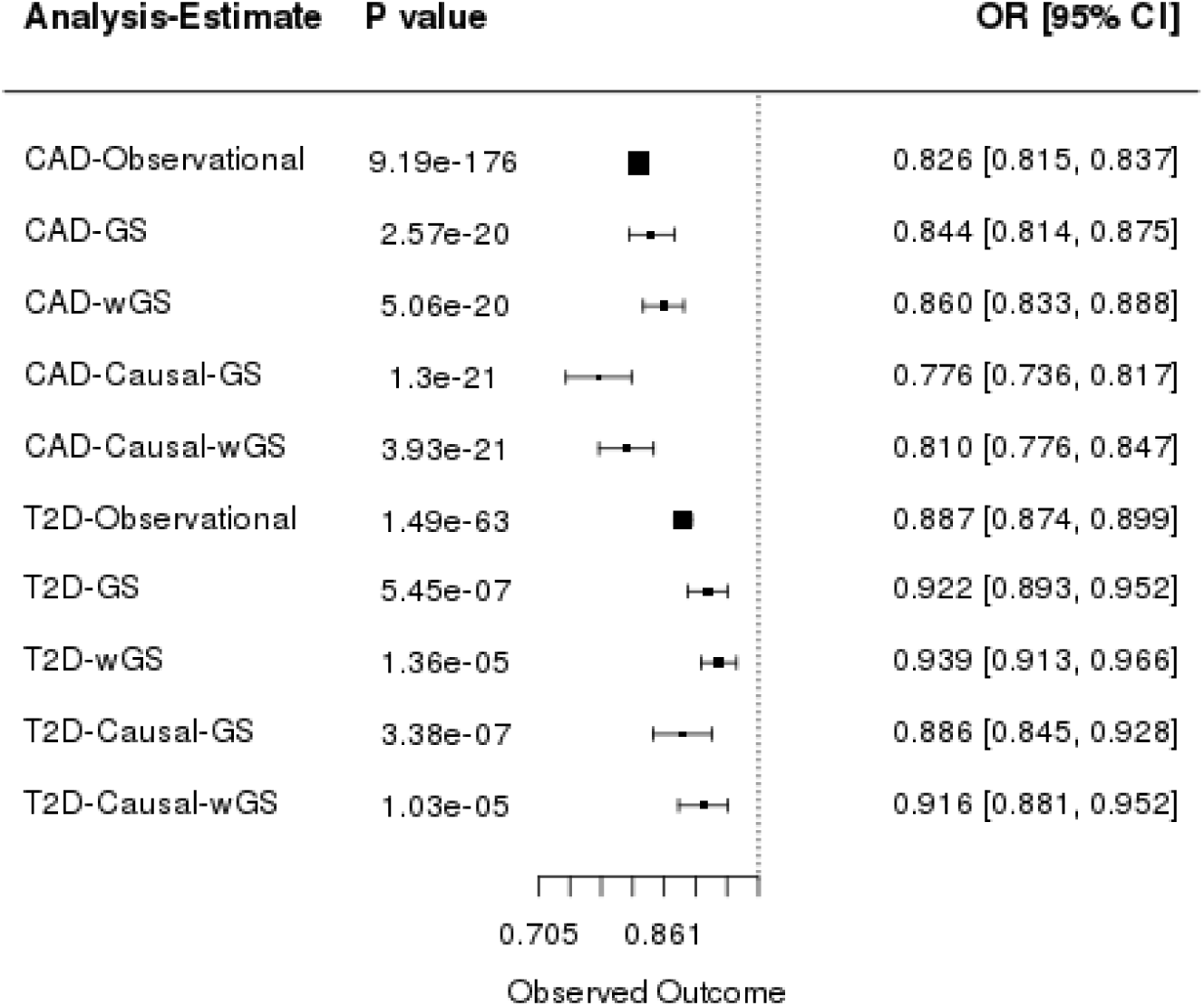
Observational and Instrumental Variables Estimates of the effect of height on cardiometabolic events. Effect estimates represent the OR (95% CI) per 1 SD increase in height, observational estimates were adjusted for age and sex. Causal estimates were derived from instrumental variable (IV) analysis.

We also performed for both CAD and T2D two-stage analysis adjusted for one risk factor at the time (BMI, SBP, DBP, hypercholesterolemia). Our results (STable 4b) suggest that the effect of height on CAD is independent of each risk factor tested whereas its effect on T2D was completely abolished after adjustment for BMI..

#### Sensitivity Analyses

We undertook a series of sensitivity analyses to investigate the causal effect between height and CAD by IVA after sequentially excluding variants nominally associated with BMI, BP or lipids (*P*<0.05 were excluded). In each case the remaining variants constitute valid instruments and their causal effect estimates will therefore be immune to confounding. After excluding the variants associated with BMI, a 1 SD increase in height, measured by the GS (wGS gave very similar results), was associated with 22% lower risk of CAD (OR= 0.78, 95% CI= 0.73 to 0.83), the same as the observed effect without instrument exclusions. Similar results were obtained also after excluding variants associated with any lipid trait, a 1 SD higher height was associated with a 13% decrease in the odds of CAD (OR= 0.87, 95% CI = 95% 0.78 to 0.88), whereas exclusion of variants associated with BP resulted in an 11% lower risk (OR= 0.89, 95% CI= 0.82 to 0.96) of CAD (STable 5, SFigure 3).

Sensitivity analyses to investigate the causal effect of height on T2D (1SD increase in height was associated with an OR of 0.88) resulted in an attenuation of the height effect after removing BMI, lipid or BP associated variants from the two-stage analysis (STable 5,SFigure 3).

For both CAD and T2D, two-stage analysis adjusted for one risk factor at the time (BMI, SBP, DBP, hypercholesterolemia) gave similar results to those described above (STable 4b). These analyses estimate the causal effect of height that is independent of each risk factor and directly estimate only the risk factor independent effect.

### Two-sample Mendelian Randomisation analyses

To further investigate the causal relationships found using two-stage analysis in UKBB and also test the validity of the GS as an instrument, two-sample MR approaches were used to detect and accommodate violations of the MR assumptions, specifically horizontal pleiotropy.

We accessed summary statistics from the largest genetic studies publically available for height (up to 700,000 individuals), CAD (up to 71,000 cases), and T2D (up to 27,000 individuals). Two-sample MR analyses were performed using the IVW method ^13^, alongside four other methods to overcome the violations of specific IV assumptions, as no single method controls for all statistical properties that may impact MR estimates - namely: WM, Egger, GSMR ^14^ and MBE ^11^ methods. Consistency in results across methods builds confidence in the obtained estimates, as they rely in different assumptions and models of horizontal pleiotropy. Funnel plots were also assessed for any deviations which can be suggestive of pleiotropy. We note that the plots appear generally symmetrical, suggesting no evidence for horizontal pleiotropy (SFigures 4b,5b).

IVW indicated a causal effect of increased height lowering CAD risk (Figure 3) consistent in direction with the IVA analyses. There was little evidence of heterogeneity in the analysis (*p*=0.9). The slope from the Egger regression was consistent with these findings (OR of 0.86 per 1 SD higher height, 95% CI= 0.79 to 0.94), and no evidence of directional pleiotropy (Intercept= −0.0009, 95% CI= −0.0029 to 0.0012) (Figure 3, SFigures 4, 5). We measured a low dilution bias in the MR-Egger casual effect, 97.5% through the I^2^ index of gene-exposure estimates, suggesting no violation of the NOME assumption (see methods) (STable 6). The results obtained using the WM approach further confirmed the direction and magnitude of effect seen with the other methods (OR= 0.83, 95% CI= 0.81 to 0.85), providing no evidence for pleiotropy (STable 6, Figure 3). The MBE method, which relaxes the instrumental variable assumptions and presents less bias and lower type-I error rates than the other methods, gave similar results; 1 SD higher height was associated with 18% decrease in the odds of CAD with the weighted method assuming the NOME assumption is valid (OR=0.82, 95%CI=0.73 to 0.92), setting the bandwidth tuning parameter φ equal to 1. STable 7. Finally, the GSMR method suggested that 1 SD higher height was associated with 16% decrease in the risk of CAD (OR=0.84, 95% CI=0.82 to 0.88, *P*= 2.91×10^-21^) (STable 8), slightly higher than the estimate from a previous study ^14^.

**Figure 3:**
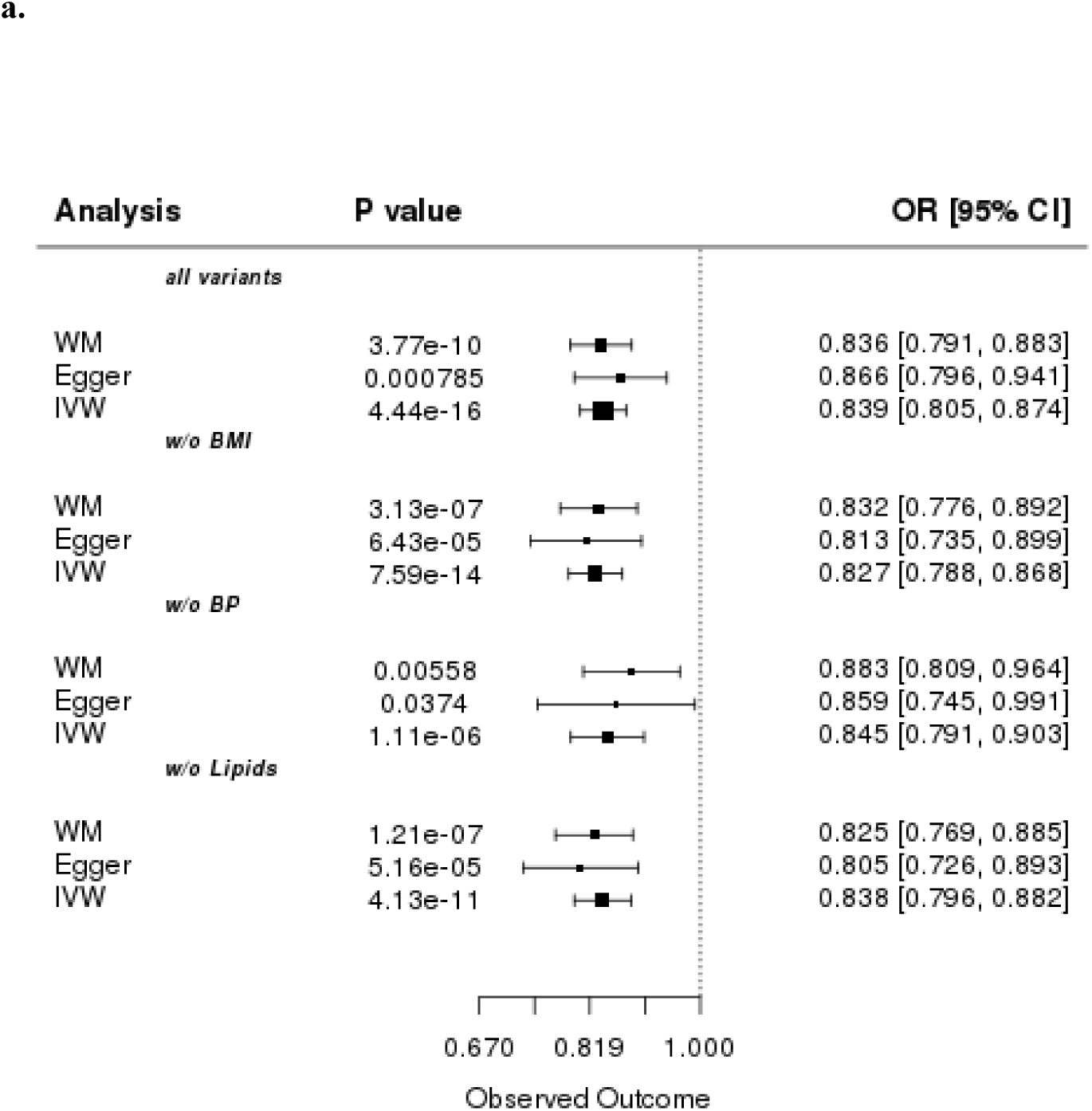
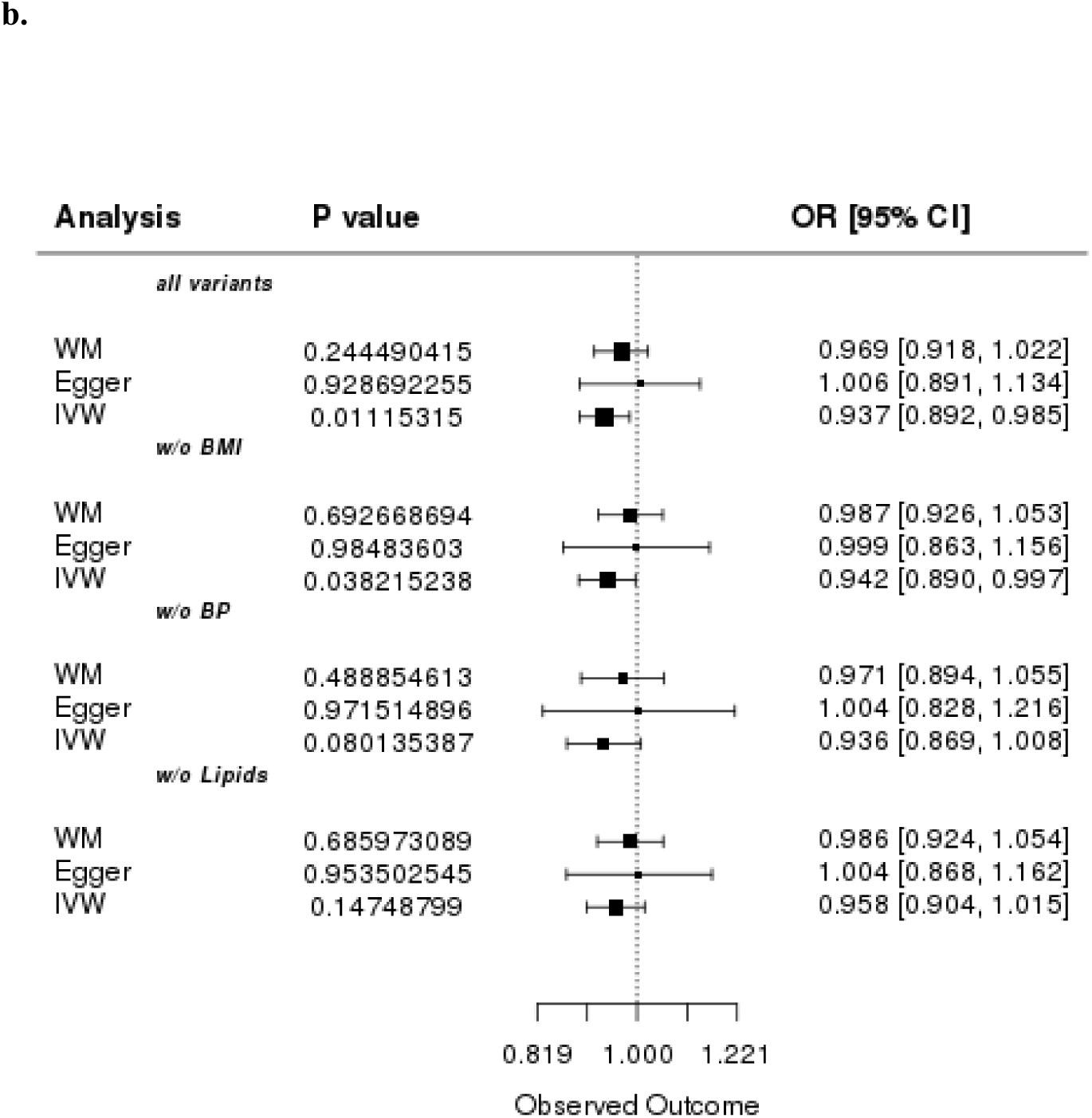
Two sample Mendelian Randomisation analyses and causal estimates from UK Biobank. Estimates of the Effect of height on A. Coronary Artery Disease and B. Type 2 Diabetes, after removing variants nominally associated with BMI, lipids or blood pressure. Effect estimates represent the ORs (95% CI)

The results we obtained for CAD using two-sample MR analyses were largely concordant with the two-stage analysis (Methods); 1 SD increase in height (6.4 cm) was associated with a 14% (OR= 0.86) lower risk of CAD with no evidence for directional pleiotropy (Egger method) (STable 6).

Sensitivity analyses showed that exclusion of variants associated with either BMI, BP, or lipid levels (i.e. one trait at a time) slightly increased the signal, a 1 SD higher height (cm) was associated with a 19% (BMI variants), 14% (-BP) or 19% (-lipids) lower risk of CAD (Stables 9-14, Figure 3A). After exclusion of any height variant nominally associated with any of BMI, lipids or BP, the remaining 155 height associated variants yielded the strongest effect with 1 SD increase in height associated with 21% lower risk of CAD (OR= 0.79, 95% CI= 0.64 to 0.97, *P*= 0.02; Stable 15).

The IVW method indicated a nominally significant causal association between height and T2D (OR=0.93, 95%CI=0.89 to 0.98) and there was no evidence of directional pleiotropy (Intercept= −0.00206, 95% CI= −0.00529 to 0.0011) (STable 16, Figure 3B, SFigure 5). Exclusion of variants associated with BMI resulted in a nominally significant causal effect of height on T2D risk (*P*=0.04), but not when we excluded variants associated with lipids (*P*=0.14) or BP (*P*=0.08) (STables 17-20, Figure 3B).Consistent with our results from the two-stage analysis in UKBB, IVW analysis when T2D was adjusted for BMI showed no causal effect of height on T2D (*P*=0.95) (STable 21).Neither the mode-based nor the GSMR method provided any evidence for a causal effect between height and T2D (STables 8, 22).

### Mediation Analyses

As described above, BMI adjustment in the MR analyses showed complete attenuation of the causal effect of height on T2D and a modest decrease of the effect on CAD. Also, when we performed sensitivity analyses i,.e. by excluding all BMI associated variants the results were unchanged. So, the MR assumption of no correlation with potential confounders (ie BMI) is thus fulfilled. Therefore, we explored BMI as potential mediator in the relation between height and CAD or T2D. Valid instruments for height and BMI were included. We applied a multiple-stage approach (see methods) in UKBB to assess the direct genetic effect of height on CAD and T2D (STable 23a). The causal effect of height on CAD after adjustment for genetic BMI was still significant (OR= 0.83, 95% CI= 0.79 to 0.87, *P*=2.79×10^−12^). However, for T2D no causal effect was observed in the multiple stage least square approach after adjustment for BMI (OR= 0.98, 95% CI= 0.93 to 1.03, *P*=0.44) (STable 23a). In addition, we performed sensitivity analyses in order to investigate the robustness of the estimation in the above mediation analysis, by excluding any height associated variants which had a nominal association with BMI. This exclusion did not affect the previous observations (CAD: OR= 0.86, 95% CI= 0.78 to 0.88, *P*=8.77×10^−10^), T2D: OR= 0.96, 95% CI= 0.91 to 1.01, *P*=0.17)

To increase power, we further investigated by Multivariable MR analysis (see Methods) the above mediation effects using summary statistics data, in order to interrogate whether the exposure is causally associated with the outcome given the risk factors. Using this approach, we estimated the effect of height on CAD and T2D risk taking account of genetically determined BMI, that is adjusting for the effect of each instrument with BMI ^15^. Using summary data we also adjusted for genetic associations with lipid levels, BP, and glycaemic traits. Multivariable MR analyses and estimation of the direct and total effect after adjusting for the effect of each possible mediator, showed that the causal effect of height on the risk of CAD was independent of the potential mediators. More specifically, the height-CAD effect reduced from 0.83 (95%CI= 0.80 to 0.87) to 0.86 (95%CI = 0.82 to 0.90) with adjustment for the estimated effects of 2h Glu (Stables 24a, and 25a, Figure 4a). The causal effect of height on T2D completely disappeared after adjusting for the genetic effect of BMI STable 24b). For T2D adjusted for BMI, there was no evidence of a direct effect of height (OR= 0.99, 95% CI= 0.96, 1.02, *P*=0.95). These observations for T2D adjusted for BMI did not change after taking into account the genetic effect of glycaemic, BP and lipid traits in Multivariable MR (STables 24b, 25c, Figure 4b). To further evaluate the robustness of the estimation in the mediation analyses, we performed the previous analyses by excluding variants nominally associated with BMI. The causal effect of height on T2D after excluding BMI associated variants also disappears after taking into account the genetic effect of BMI (STable 25d).

**Figure 4:**
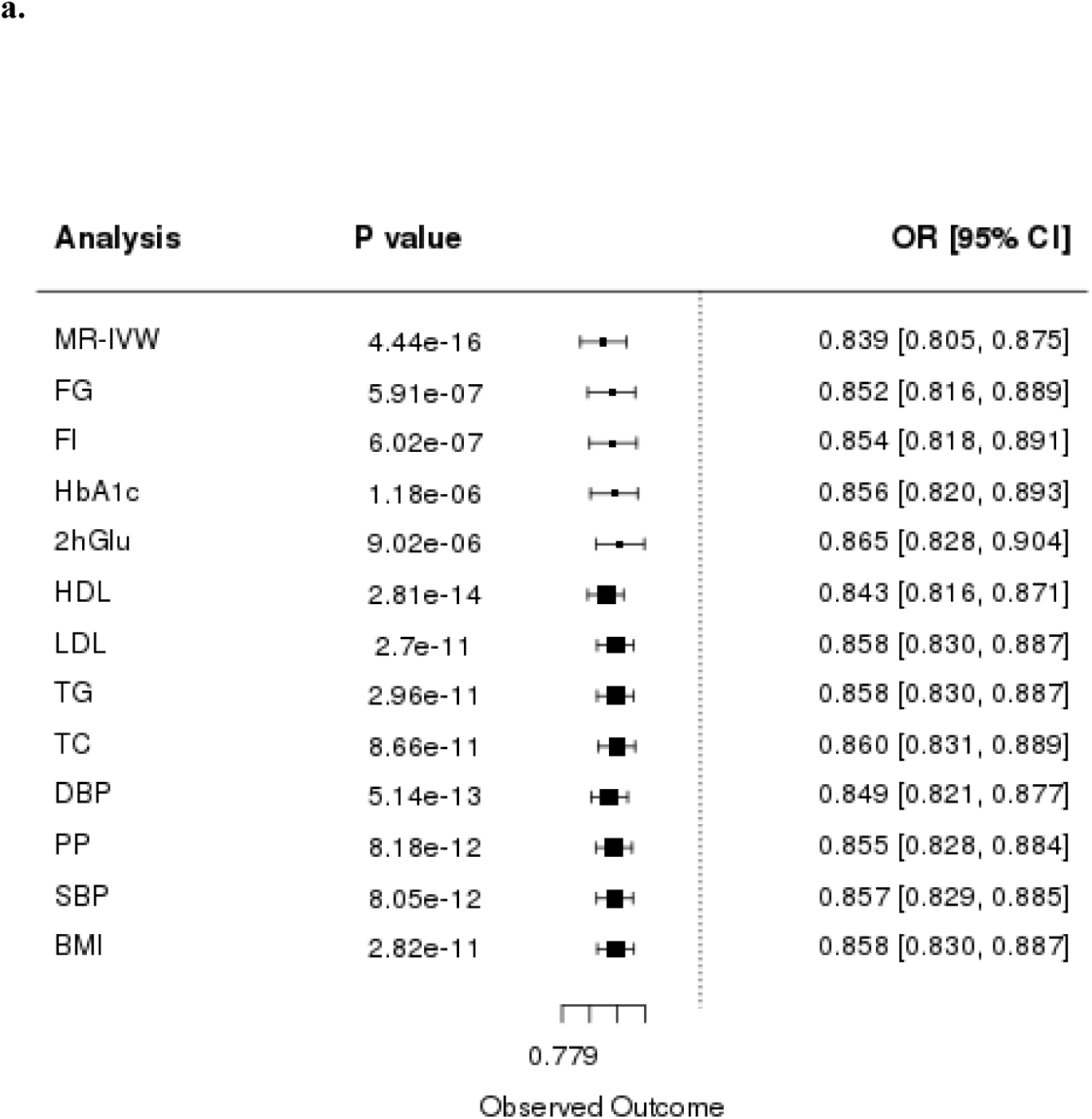
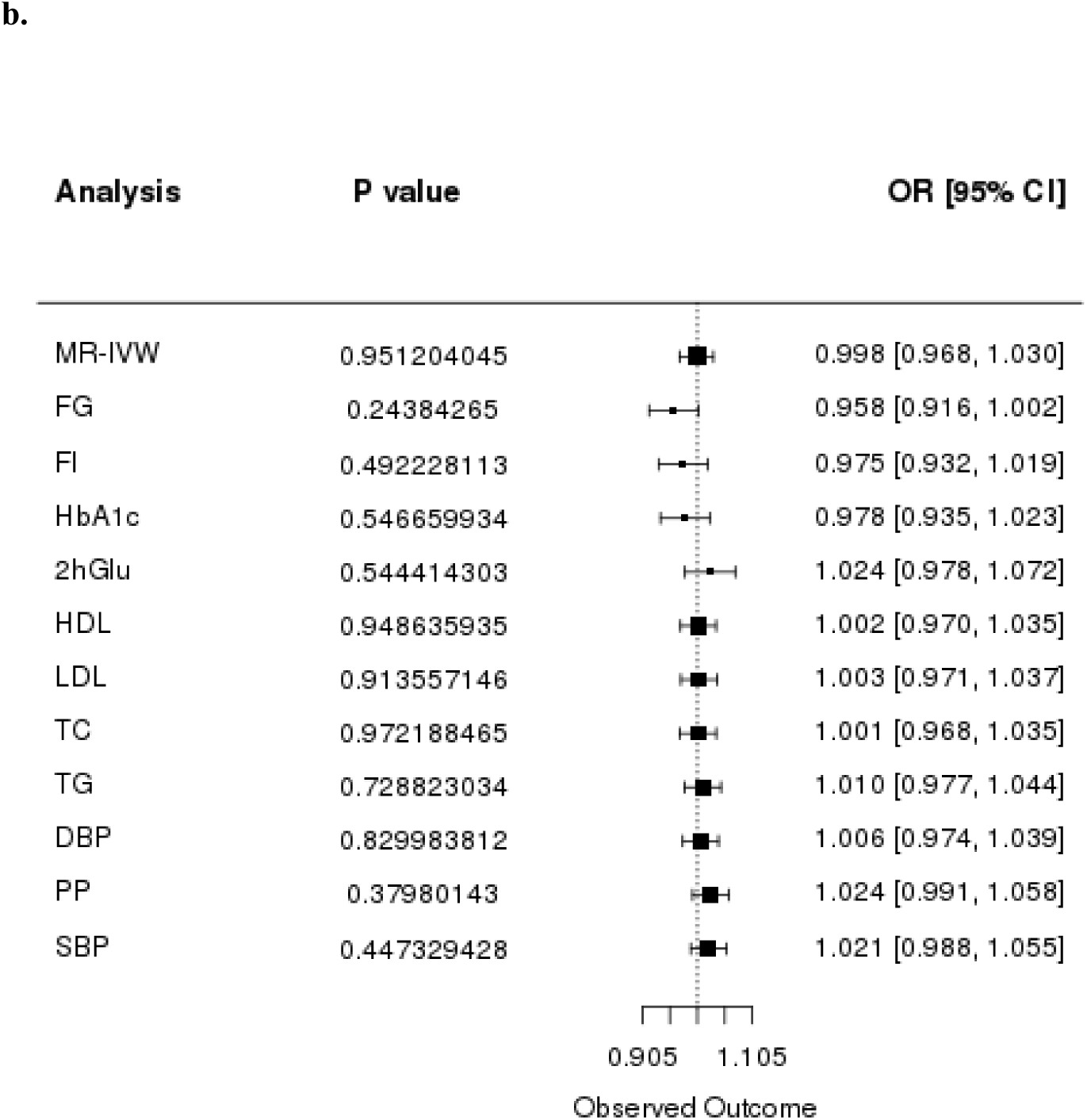
Multivariable separate-sample MR analysis of the effect of height (per SD) on A. CAD and B. T2D. Analysis indicates the genetic effect of each trait.

### Bidirectional Mendelian Randomisation analysis

As a negative control, we investigated the effect of the known CAD associated variants (from CARDIOGRAMplusC4D) ^16^ and T2D (from DIAGRAM consortium) ^17^ on adult height by two-sample MR using summary statistics data. The corresponding association statistics of these variants with height were extracted from UKBB (STables 28, 29).

When we performed the MR analyses using variants related to T2D as instruments for height, there was no evidence of a causal effect of T2D on height (*P*=0.39) and no evidence or directional pleiotropy (*P*=0.62) from the MR-Egger regression (STable 26).

The analysis using genetic variants related to CAD as instruments for height measurements, indicating no evidence for a causal effect of CAD (*P*=0.57) on height (STable 27).

## Discussion

To the best of our knowledge this is the first study investigating not only the causal relationship between adult height and cardiometabolic disease (CAD and T2D) but also the extent to which traditional risk factors (obesity, glycaemic, lipid, and BP) may mediate such effects.

Consistent with previous studies^4, 18^, our MR results provided strong evidence for a protective causal effect of adult height on CAD risk, 1 SD higher height (~6.5 cm) was causally associated with a 22% lower risk of CAD (OR= 0.84, 95% CI= 0.73 to 0.81) in UKBB.

It has been postulated that the inverse association between height and CAD may be due to shorter individuals having higher BP ^19^. Shorter individuals have increased heart rate, increased augmentation of the systolic pulse, which may increase ventricular systolic work ^20^. Furthermore, shorter individuals have smaller vessel calibre, so their arteries can become more easily occluded and subsequently increase CAD risk ^20, 21, 22, 23^.

### Potential mechanisms

#### CAD

Exclusion of variants that are nominally associated with BP only marginally attenuated the causal association between height and CAD (OR 0.78 vs 0.89; STable 6). Similar results were obtained in analyses adjusting for BP traits and Multivariable MR (STable 7). Lipids appeared to have an even more modest effect than BP on the causal effect of adult height on CAD risk (STables 6 and 7) in the different MR approaches.These results suggest that the effect of height in cardiometabolic disease involve alternative biological pathways. However, one caveat in these analyses is that adult height is known to be associated to socio-economic factors that may also mediate the observed effect ^24^.

#### T2D

In contrast to our findings regarding CAD, we did not find any evidence supportive of a causal effect of height on T2D after adjustment for BMI, despite the modest association we observed based on observational data. Indeed, our mediation analyses suggest that the causal effect of height on T2D is mediated by BMI.

### Strengths and limitations

There was concordance between the causal instrumental variable analysis and the minimally adjusted observational analyses for the cardiometabolic diseases we interrogated. This indicates that there is not extensive unmeasured confounding in the observational estimates. This is not surprising since height is extremely heritable (little possibility for non-genetic confounding) and established decades before onset of the examined diseases.

Our results suggest that the observed relationships of height with the CAD are very close to the unidirectional causal ones and are less susceptible to arise from bias or confounding. A potential limitation of our study is that we have assumed no interaction between height and the mediators, but we were unable to test for this as we used aggregated genome-wide data for glycaemic and lipid traits (unavailable in UK Biobank at the time of analyses). Also, in the two-sample MR, where independent samples were used, weak instrument bias may result in bias towards the null. Similarly, in mediation analyses weak instrument bias could result in underestimation of the mediating effects. However, we assumed that estimates come from two different homogenous population studies without overlapping samples. The use of large sample sizes and instruments with large F statistics in our analyses is likely to have minimised any effect on the obtained results. Also, the IVW method has been shown to lead to slightly biased estimates (10% in either direction) in the presence of binary outcome and to a natural correlation between causal estimates and standard errors that could contribute to the presence of heterogeneity misinterpretable as pleiotropy ^25^.

In summary, we show that height has a causal relation with CAD, and this association is still present after taking into account the genetic effect of possible mediators including lipid, glycaemic, BP and anthropometric traits. We also show that BMI mediates the effect of height on T2D risk. Our findings may support the need for interventional studies to examine whether lowering BMI in individuals with defective growth is beneficial at reducing their risk of T2D. Similarly, lifestyle changes that may boost healthy nutrition and skeletal growth in the population may be advantageous towards the risk of CAD later in life.

## Online Methods

### Analyses using individual level data

The UKBB recruited more than 500,000 individuals aged 37-73 across the country between 2006 and 2010. All participants provided information with questionnaires and interviews regarding health status, anthropometric characteristics, blood, urine and saliva samples^10^. Data underwent central quality control (see (http://biobank.ctsu.ox.ac.uk). UKBB samples were excluded due to sample relatedness determined as kinship coefficient greater than 0.0884.

Height (cm) was measured using a Seca 202 device in all participants of UKBB. We excluded individuals who exceeded a +-5 SD away from the mean of the sampled population.

BMI was constructed from height and weight measured during the initial Assessment Centre visit. Value is not present if either of these readings were omitted.

Continuous traits were adjusted for demographics, genetic structure and converted to a normal distribution to limit the influence of any population stratification and provide standard deviation effect sizes. Residuals of the exposure from standard linear regression were taken by using as covariates: age, sex, five principal components and batch. The residuals were then inverse normalized in order to improve comparability with summary data MR analysis.

#### CAD definitions

UKBB self-reported data: ‘Vascular/heart problems diagnosed by doctor’ or ‘Noncancer illnesses that self-reported as angina or heart attack’. Self-reported surgery defined as either PTCA, CABG or triple heart bypass. HESIN hospital episodes data and death registry data using diagnosis and operation -primary and secondary cause: MI defined as hospital admission or cause of death due to ICD9 410-412, ICD10 I21-I24, I25.2; PTCA is defined as hospital admission for PTCA (OPCS-4 K49, K50.1, K75); CABG is defined as hospital admission for CABG (OPCS-4 K40 –K46); Angina or chronic IHD defined as hospital admission or death due to ICD9 413, 414.0, 414.8, 414.9, ICD10 I20, I25.1, I25.5-I25.9.

#### Type 2 Diabetes definitions

UKBB self-reported data: ‘Diabetes by Doctor’ or ‘‘Non-cancer illnesses that self-reported as T2D’. HESIN hospital episodes data and death registry data using diagnosis and operation -primary and secondary cause: T2D defined as hospital admission or cause of death due to ICD10 E11.

#### Hypercholesterolemia definitions

UKBB self-reported data: ‘Non-cancer illnesses that self-reported as Hypercholesterolemia. HESIN hospital episodes data and death registry data using diagnosis and operation -primary and secondary cause: Hypercholesterolemia defined as hospital admission or cause of death due to ICD10 E780, E7800, E7801.

We performed conventional regression analysis of each cardiometabolic disease (CAD and T2D) against height by using logistic regression. Height was the independent variable for each trait of interest by using linear and logistic regression for continuous and binary traits, respectively. These associations were adjusted for age, sex, first 5 PCs and batch. This information was compared with the estimates derived from instrumental variable analyses.

Genotypes were extracted from UKBB imputation dataset (STable 30: Summary of the height variants previously identified as associated with height at genome wide significance). Individual genotypes were excluded if the imputation quality was less than 0.4. We confirmed that the variants were imputed with high quality by comparing them with the directly genotyped data, where available. 829 independent (r^2^ ≤0.05) and GWA significant (p<5×10^-8^) SNPs were selected from the largest GWA studies for height ^6, 7^. One condition of MR is that exposure-related SNPs (the instrumental variables) must not be in LD with each other, as that can result in confounding ^3^. For this purpose LD between all variants was estimated in European samples from 1000 Genomes using Plink software version 1.9 ^26^. When two or more SNPs were in LD (r^2^ >0.05) only the most strongly associated variant with height, based on p-value, was kept. Two genetic scores, weighted and unweighted, were created. The first incorporated 829 independent height associated variants LD pruned at r^2^ of 0.05, which were associated with height at genome-wide significance in the GIANT studies of up to 700,000 individuals (STable 22). Variants with low imputation quality or unavailable were excluded. Individual variants were recoded as 0, 1 and 2, depending on the number of height increasing alleles. These variants were used to create genetic scores.

The unweighted genetic scores (GS) for each individual were created by summing the number of height increasing alleles for the 829 SNPs they are carrying. Weighted genetic scores (wGS) were also modelled. The wGS was calculated as the sum of the number of height-associated alleles, weighted by the relative effect size (β coefficient) reported from the discovery meta-analysis ^6, 7^. In the derived wGS, β represents the association between an additional weighted height-associated allele at each single nucleotide polymorphisms (SNP) and height: weighted score=β_1_×SNP_1_+β_2_×SNP2+…β_n_×SNP_n_. We present the range of the possible number of weighted height-increasing alleles, by dividing the score by the average effect size of the variants for each individual ^27^. This is a transformation of the wGRS so that the range equaled that of the unweighted score. Linear regression for each score with height and logistic regression for each score with disease status were performed.

### Mendelian randomization

SNPs from the largest GWAS study of height to date were identified by the 2015 and 2017 summary statistics files from the GIANT (Genetic Investigation of Anthropometric Traits) consortium. Data on effect and other alleles for each of the 829 LD pruned variants in up to 700,000 individuals of European descent, along with allele frequencies, beta coefficients for allele dose, and a 6.4 cm change in height, p-value and standard errors (SE) were extracted. |In order to test the statistical significance of the association of the instrument with height, an F statistic was calculated using the formula: (β exposure ×β exposure)/(se exposure × se exposure) ^28^.

We first performed an instrumental variable analysis -IVA (two-stage analysis) in UKBB, where we had access to individual level data, and then expanded this analysis to the largest summary statistics data sets currently available, assuming homogeneity among the studies. We used data from genetic studies on height (GIANT) ^6, 7^, CAD (CARDIoGRAMplusC4D) ^16^, T2D (DIAGRAM)^17^ using two-sample MR methods. To investigate potential mediators, genetic associations with (i) fasting insulin, fasting glucose, 2hGlu and HbA1c were obtained from MAGIC, http://www.magicinvestigators.org/). (ii) HDL-cholesterol, LDL-cholesterol, total cholesterol and triglycerides were obtained from GLGC (http://csg.sph.umich.edu/abecasis/public/lipids2013/), (iii) anthropometric traits for GIANT (https://portals.broadinstitute.org/collaboration/giant/index.php/GIANT_consortium_data_files) and (iv) BP from ICBP ^29^.

MR relies on certain assumptions (Figure 5):

**Figure 5:**
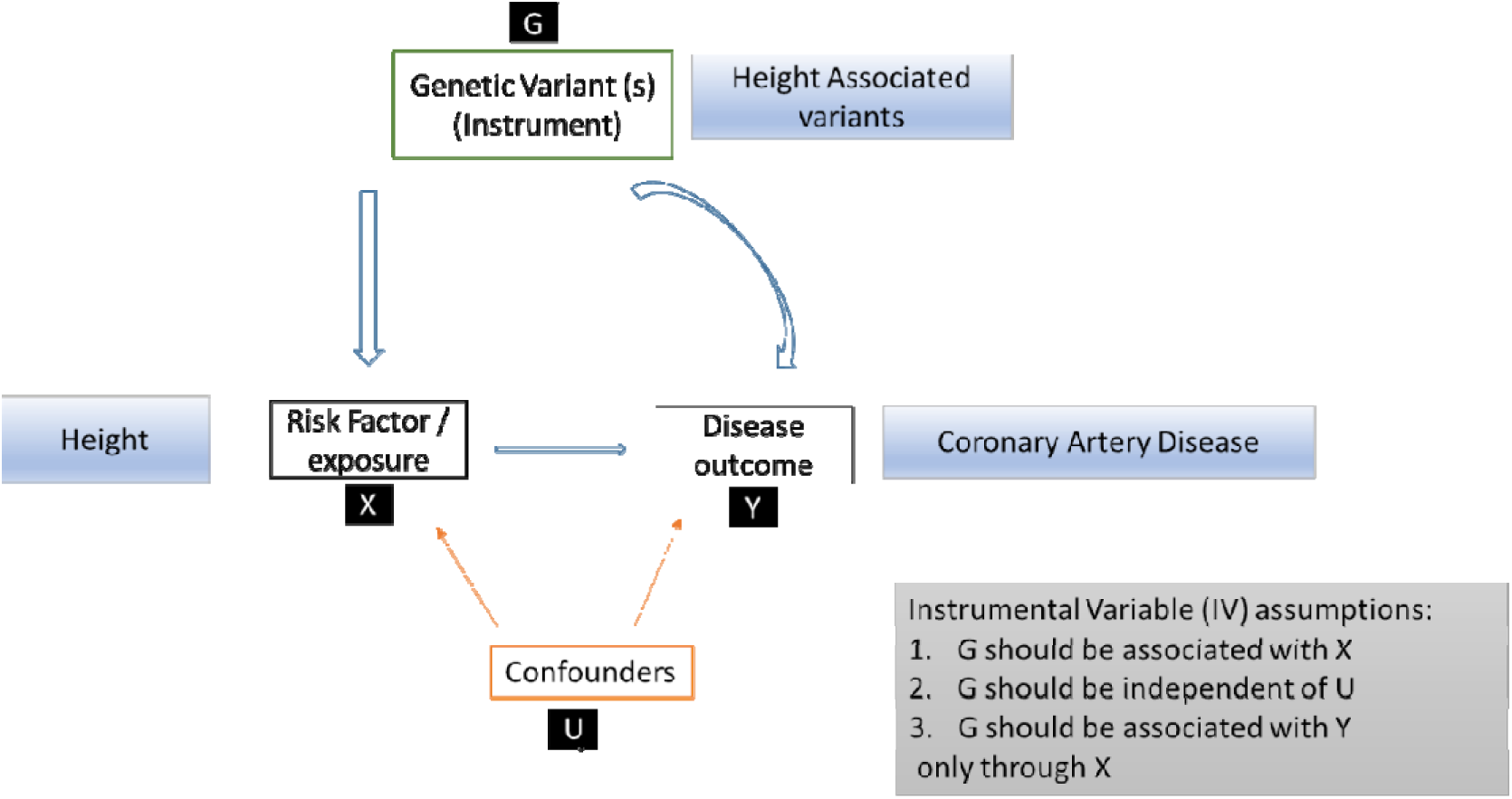
Mendelian Randomisation Study Design

1. The IV is robustly associated with the exposure of interest. This can be evaluated by estimating the F statistic and the R^2^ value. It is substantial to have large studies, especially in instances where the IV explains a small amount of the variance in the exposure (R^2^). Genome wide association studies for height have yielded a large number of genetic variants that account for around 30% of height heritability. That allows the use of strong instruments to be developed.
2. The IV has to be independent of any confounder. When using individual level data, known confounders can be checked. In two-sample MR, confounders can obstruct testing of this assumption due to lack of summary data results on the association between the candidate genetic instruments and the confounders.
3. The IV is independent of the outcome, given the exposure and any possible confounders.

The IV should not influence the outcome on an alternative path, other than through the exposure. This assumption is violated by horizontal pleiotropy, in which there are alternative pathways that the IV can affect the outcome.

#### Instrumental Variable Analysis (Two-stage analysis)

The MR approach used in this study was based on the following assumptions: the height genetic scores had a strong association with measured height; the height genetic scores were not associated with confounding factors that could bias the observational association between height and cardiometabolic disease; the height score was related to the outcome only through its effect on the exposure, assuming a linear relationship between height and the logit-transformed outcome.

In order to estimate the causal effect of height on disease status we performed instrumental variable analysis by using height genetic score as instrument. For the binary traits, we used the two-stage estimator (logistic control function estimator) ^30, 31, 32^. The analysis was performed in two stages. First, the association between height genetic score and height was assessed. These predicted values were then used as the independent variable and disease status as the dependent variable in a logistic regression model. Analyses were adjusted for age, sex, first five PCs and batch.

#### Two-sample Mendelian Randomisation

Two-sample MR was undertaken using GWA summary data from separate samples, where data of the genotypes and the exposure of interest are available in one sample, and data on genotype and the outcome of interest are available in the other. For this part no ethical approval was sought as all data were derived from summary statistics of published GWAS studies, with no individual-level data used.

#### Association of height variants with cardiometabolic traits

Coronary artery disease genotyping data were derived from the most recent meta-analysis of Cardiogram+C4D, which investigated the association of 7M variants after imputation in up to 30,000 cases. The per-allele log-OR of CAD was extracted together with its SE for each of the independent genome-wide significant height variants. Effect sizes were aligned to the height increasing allele.

The two-sample MR was undertaken using previously described methods^12^. Wald ratios were estimated for each SNP by dividing the per allele log-OR for CAD (beta_gy) by the per-allele effect on height for each SNP (beta_gx). SE for each Wald ratio was derived from the SE of the variant-outcome association divided by the variant-exposure association for each instrument. We calculated the Wald ratio estimate where outcome ~ GS and exposure ~GS estimates were obtained using the previous regression models with the GS.

#### Inverse-variance weighted (IVW) method

Conventional linear regression analysis of the variant-exposure association and variant-outcome association for each instrument was undertaken and weighted by inverse variance. The point estimate is equal to that derived from fixed-effect meta-analysis. The IVW method assumes that all variants are valid IVs. An IVW corrected for the standard errors of each instrument, for the correlation between the associations of the instrument with the exposure and the association of the instrument with the outcome. When the IVW method shows substantial heterogeneity, this means that there may exist alternative pathways through which the SNPs affect the outcome (horizontal pleiotropy). Heterogeneity for the Wald ratios was tested with the Cochran’s Q and quantified with the I^2^ index^33^.

#### MR-Egger Method

MR-Egger method is more robust to potential violations of the standard instrumental variable assumptions. This method was used to address the issue of the aggregate unbalanced horizontal pleiotropy, which could violate the third assumption of IV analysis. MR-Egger is similar to the IVW method except that the intercept is not constrained to pass through the origin ^12^. MR-Egger method uses a weighted regression with an unconstrained intercept to regress the effect sizes of the variant-outcome associations against effect sizes of variant-exposure associations. The unconstrained intercept removes the assumption that al genetic variants are valid instruments. This method is less prone to confounding from possibly pleiotropic variants which could have stronger effects on outcomes compared to the effect on the exposure.

When a non-zero intercept from the MR-Egger is observed, that would suggest that there are pleiotropic effects; this could result in bias of the IVW estimates, which are in the direction indicated by the intercept term.. The estimate for the effect of the exposure on the outcome, is provided by the slope of the MR-Egger. It is important to mention that this estimate is correct, taking into account an additional assumption, the InSIDE (instrument strength independent of direct effect) assumption, which means that the associations between the genetic variants and the exposure are independent of the effect that he variants have directly on the outcome. Funnel plots can be used to demonstrate the individual variant effects on the exposure and the outcome against the inverse of their SE. When no pleiotropy is present, the IV estimates for each variants are symmetrically distributed around the point estimate. However, it is possible to understand the contribution of each IV on the overall Q statistic graphically.

IVW and Egger methods use weights that consider the SNP-exposure associations to be known rather than estimated. This is called as the “NO Measurement Error” assumption (NOME). We use an adaptation of the I^2^ statistic in order to quantify the strength of NOME violation for MR-Egger. This measure is called IGX2 and lies between 0 and 1. A high value of IGX2 and close to 1, indicates that dilution does not affect the standard MR-Egger analyses performed ^33, 34^. For the main analyses we used the first order weights, which correspond to the first term of the Taylor series expansion, which approximate the standard error of the Wald ratio estimate ^34^. The IVW and the MR-Egger regression analyses were repeated using the second order weights, which correspond to the first two terms of the Taylor series expansion ^33^, doi: https://doi.org/10.1101/159442.

#### Weighted Median method

The weighted median method was used to investigate pleiotropy ^35^. In this method, the MR estimates are ordered and weighted by the inverse of their variance. When more than 50% of the total weight comes from SNPs without pleiotropic effects, the median MR estimate should remain unbiased. This method improves precision and is more robust to violations of the InSIDE assumption. If InSIDE holds, then MR-Egger is consistent, while the weighted median will be only if 50% of the total weight comes from SNPs without pleiotropic effects.

#### Mode-based estimate (MBE)

The MBE method has presented less bias and type-I error rates in simulations than other methods under the null in many situations. The MBE relaxes the instrumental variable assumptions and is less prone to bias due to violations of the InSIDE assumption^11^.

**GSMR** (Generalised Summary-data-based Mendelian randomisation)

Given that the correlations between the SNP instruments could lead to biased (smaller) MR standard errors ^13^, we also applied a method (GSMR) that performs a multi-SNP Mendelian Randomisation analysis using GWAS summary-level data accounting for the sampling variance in the estimated SNP effects and remaining LD between SNPs. The gsmr R-package implements the GSMR method to test for putative causal association between a risk factor and a disease. We used 1000 Genomes (European population) imputed data to create the LD correlation matrix ^14^.

### Sensitivity Analyses to investigate the presence of pleiotropy of Height on Coronary Artery Disease and type 2 Diabetes

To further investigate the presence of pleiotropy and narrow down the set of height variants which may have a causal effect on the risk of CAD and T2D, we performed sensitivity analyses.

First, we assessed which variants may contribute to the total heterogeneity, by estimating the Q statistic for each instrument. We then used some different thresholds (5th (L1), 1st (L2), 0.19th (L3) percentile of a chi-squared with 1 degree of freedom) and excluded variants which had a Q>L3, Q>L2 and Q>L1. Second, we assessed which variants are associated with any potential mediators including BMI, BP (SBP, DBP, PP) and lipids (LDL, HDL, TG, TC). We excluded any height associated variant which showed evidence for association (nominally significant) with these traits.

### Mediation Analyses

To estimate the effect of height on T2D and CAD taking into account the role of potential mediators, we performed multivariable MR analyses, by using the IVW MR method with summary statistics data, after adjusting for the effect of each instrument with the potential mediator ^36^. We evaluated the proportion of the effect that is mediated by any of the potential mediators by the changes in the total effect of the genetically determined height and on the outcomes, assuming that the mediators are continuously measured variables (Multivariable MR). We estimated the total, direct and indirect effects of the risk factor on the outcome by using summary data ^15^. It is recommended to provide estimates of the total and direct effects, but not the indirect effect, as calculation of the indirect effect relies on the linearity of the relationship that cannot occur with a binary outcome ^15^.

Using individual level data we also applied a three stage approach where we first estimated the fitted values of the height genetic risk score with height, second the fitted values of a BMI genetic risk score with BMI and finally we used these fitted values to estimate the direct effect of height on the outcomes ^32, 37^. The weighted regression method for calculating the direct effect is also equivalent to a two-stage regression method, except that the first stage also regresses the mediator on the genetic variants, and the second stage regresses the outcome on fitted values of the exposure and fitted values of the mediator. This two-stage approach can be undertaken to estimate the direct effect when individual-level data are available ^15, 32^.

### Bidirectional Mendelian Randomisation analyses

We performed bidirectional Mendelian Randomisation analyses of the association of Coronary artery disease and height. To construct a genetic instrument for CAD, we used variants which reached genome-wide significance in the CARDIOGRAM+C4D consortium. For the SNP-exposure we used the effect estimates and SE of the associations of each variant with CAD derived from the meta-analysis^16^. For the SNP -outcome associations we used effect estimates and SE of the association of the CAD – related variants with height from the UKBB (STable 29). We used a similar approach for the bidirectional MR analyses with T2D. We used genome-wide associated with T2D in the DIAGRAM consortium ^17^ (STable 30). Two-sample MR analyses were conducted as described above for height and coronary artery disease. All statistical analysis was performed using R (version 3.4.3, the R Foundation for Statistical Computing, Vienna, Austria) software.

## Abbreviations

GIANT: Genetic Investigation of Anthropometric Traits
CARDIoGRAM: Coronary ARtery DIsease Genome wide Replication And Meta-analysis
DIAGRAM: DIAbetes Genetics Replication And Meta-analysis
GLGC: Global lipids genetics consortium
MAGIC: Meta-analyses of glucose and insulin related traits consortium
ICBP: International Consortium for Blood Pressure

## Acknowledgments

E.M. is supported by the British Heart Foundation (BHF) grant RG/14/5/30893 to P.D. Data on coronary artery disease have been contributed by the CARDIoGRAMplusC4D and UK Biobank CardioMetabolic Consortium CHD working group who used the UK Biobank Resource (application number 9922).

The work of H.R.W., M.J.C., and P.D. is supported by the NIHR Biomedical Research Centre at Barts. M.J.C. is an NIHR Senior Investigator.

CM-G: is supported by the Netherlands Organization for Health Research and Development (ZonMw VIDI 016.136.367)

## Footnotes

### Author Contributions

Study conception: E.M., F.D.G.M., C.A, J.N.H, R.J.F.L., Z.K., P.D.

Data analysis: E.M., F.D.G.M., J.Y., Z.Z, Z.K.

Writing of the manuscript: E.M., F.D.G.M., J.N.H, R.J.F.L., Z.K., P.D.

Interpretation of the results and critical revision of the manuscript: E.M., F.D.G.M.,C.A., J.Y., S.A., S.B., M.J.C., E.E., B. M., C.M.G., J.V.O., H.W., Z.Z., J.N.H, R.J.F.L., Z.K., P.D.

